# Effects of Aging on the Encoding of Spatial Direction in the Human Brain

**DOI:** 10.1101/791228

**Authors:** Christoph Koch, Shu-Chen Li, Thad A. Polk, Nicolas W. Schuck

## Abstract

Human aging is characterized by impaired spatial cognition and reductions in the distinctiveness of category-specific fMRI activation patterns. Yet, little is know about age-related decline in neural distinctiveness of spatial information. Here, we asked whether neural tuning functions of walking direction are broadened in older versus younger adults. To test this idea, we developed a novel method that allowed us to investigate changes in fMRI-measured pattern similarity while participants navigated in different directions in a virtual spatial navigation task. We expected that directional tuning functions would be broader in older adults, and thus activation patterns that reflect neighboring directions would be less distinct as compared to non-adjacent directions. Because loss of distinctiveness leads to more confusions when information is read out by downstream areas, we analyzed predictions of a decoder trained on these representations and asked (1) whether decoder confusions between two directions increase proportionally to their angular similarity, (2) and how this effect may differ between age groups. Evidence for tuning-function-like signals was found in the retrosplenial complex and primary visual cortex. Significant age differences in tuning width, however, were only found in the primary visual cortex, suggesting that less precise visual information could lead to worse directional signals in older adults. Yet, age differences in visual tuning were not related to behavior. Instead, directional information encoded in RSC correlated with memory on task. These results shed new light on neural mechanisms underling age-related spatial navigation impairments and introduce a novel approach to measure tuning specificity using fMRI.

## 1 Introduction

A central goal of aging research is to understand how aging-related neurobiological changes affect computational functions of the brain. One important approach has been to investigate how aging changes the representation of sensory information in the brain (Voss et al., 2008; Carp, Park, Polk, & Park, 2011; Schmolesky, Wang, Pu, & Leventhal, 2000), which in turn might affect cognitive operations that rely on these representations (Baltes & Lindenberger, 1997; Li, Lindenberger, & Sikström, 2001). A prominent finding in this regard is that neural patterns are less specific to the category of sensory information in older adults, a phenomenon commonly referred to as *neural dedifferentiation* (e.g. D. C. Park et al., 2004; Koen & Rugg, 2019, for recent reviews). Here, we studied age-related neural dedifferentiation in the domain of spatial navigation.

In particular, in this study we asked if aging changes how brain areas sensitive to visual and spatial information encode angular walking direction during navigation (Cullen & Taube, 2017; Blair & Sharp, 1996). In young animals, electrophysiological recordings of visually- and direction-sensitive neurons in primary visual cortex (V1) (De Valois & De Valois, 1980) and the thalamus (Taube, Muller, & Ranck, 1990a, 1990b) have revealed that although most neurons have a preferred stimulus, they are not firing in an all-or-none fashion. Rather, cells tend to fire proportionally to the similarity between the observed stimulus and their preferred stimulus, exhibiting response properties that are well approximated by a so-called Gaussian ‘tuning function’ centered around the preferred stimulus. Modelling work has also shown that a population of cells with those tuning properties will optimally encode an approximately Gaussian likelihood function of the stimulus given the population response; and suggested that this likelihood function is read out, or decoded, by downstream populations that compute optimal behavior based on sensory input (Jazayeri & Movshon, 2006; Averbeck, Latham, & Pouget, 2006). The focus of the present paper was therefore to understand age-related differences in the properties of population-based tuning functions that encode directional information.

Understanding age-effects on population-level tuning properties is important given the large number of previous investigations that have suggested a loss of specificity of neural representations in older animals and humans. This originated from reports of fMRI activation patterns in inferior temporal cortex losing categorical specificity with increasing age, i.e. activity patterns evoked by face-, place- or word- stimuli are more similar in older versus younger adults (e.g. D. C. Park et al., 2004; Voss et al., 2008; Burianová, Lee, Grady, & Moscovitch, 2013; Carp et al., 2011). Neural dedifferentiation has also been linked to memory impairment with older age (Zheng et al., 2018; Koen, Hauck, & Rugg, 2019) and related changes to similarity of neural representations might play a crucial role in the encoding and retrieval of memory content (Koen, Hauck, & Rugg, 2018; Sommer et al., 2019). Moreover, electrophysiological recordings in V1 of senescent Rhesus monkeys have found that tuning curves of orientation responsive neural populations broaden with age, effectively widening the spectrum of orientation angles a single neuron responds to (Leventhal, Wang, Pu, Zhou, & Ma, 2003; Schmolesky et al., 2000).

According to the *neural broadening hypothesis* these changes in firing properties of neural populations are a potential mechanism behind neural dedifferentiation, a notion which found support in a recent fMRI study (J. Park et al., 2012).

However, while electrophysiological recordings showed broadening within a single, continuous domain (e.g. visual orientation), the fMRI evidence is based on increased pattern similarity across distinct domains processed in anatomically separate brain areas (e.g., faces vs. houses). This is an important difference because the broader tuning functions over a continuous domain found in animals likely relate to changes in local inhibitory control (Leventhal et al., 2003). The mechanisms underlying cross-category dedifferentiation across areas as found in humans, on the other hand, must be non-local and are generally much less well understood. Thus, our focus on age-related changes in tuning properties of direction sensitive areas would allow us to build a closer link to animal studies. Moreover, the investigation of representations underlying spatial navigation might lead to insights into why age-related memory impairments are particularly pronounced in the spatial domain (Moffat, 2009; Lester, Moffat, Wiener, Barnes, & Wolbers, 2017), since spatial memory relies on a sense of direction, for instance during path integration (McNaughton, Battaglia, Jensen, Moser, & Moser, 2006; Seelig & Jayaraman, 2015).

To investigate age-related changes in visual and directional encoding of angular walking direction, we analyzed fMRI data from a previous study that used a spatial virtual reality (VR) navigation paradigm (Schuck, Doeller, Polk, Lindenberger, & Li, 2015). This work has shown that the neural underpinnings of different spatial navigation strategies are changed, and partly dedifferentiated in older adults (see also: Schuck et al., 2013). In the present paper we went beyond this work by investigating the encoding of directional information that is involved in any spatial strategy. Our hypotheses were threefold: first, we expected that fMRI signals stemming from directionally- and visually-tuned neural populations will allow us to decode walking direction above chance (directional and visual similarity were linked in the present data, as they are in daily life). Second, the similarity of two representations arising from different directions should be inversely proportional to the angular difference between these directions. Because our focus was on representational structure from the perspective of downstream areas which read out population level tuning functions (Jazayeri & Movshon, 2006; Averbeck et al., 2006), we investigated the probability of a decoder in confusing similar patterns, rather than the similarity directly. A tuning function-like signal should lead to systematically more confusions between neighbouring directions, effectively taking the shape of a Gaussian tuning function as seen in the analysis of electrophysiological recordings in animals (Mazurek, Kager, & Van Hooser, 2014). Finally, our most central hypothesis was that older adults should show decreased specificity of directional representations, which we tested by comparing the width of the fMRI-derived tuning functions.

## 2 Materials and Methods

### 2.1 Participants

This study is a re-analysis of data from 26 younger (21–34) and 22 older (56–74) male participants, as reported in Schuck et al. (2015). In addition to the exclusion criteria used in the original study (insufficient task performance, signal loss), we excluded participants with an unsuitable distribution of walking direction events that resulted in too little data for at least one direction to train the classifier (three participants, one younger, two older; for details see supplementary material section one). Additionally, one younger and one older participant had to be excluded due to missing directional information or excessive motion during the task, respectively. Therefore, 43 participants (24 younger, 21–34 years, *μ*_age_ = 27.87, *σ*_age_ = 4.01; 19 older, 56–74 years, *μ*_age_ = 67, *σ*_age_ = 3.93) entered the analysis. Additional subject characteristics can be found in (Schuck et al., 2015).

### 2.2 Virtual Reality Task

Participants performed a desktop-based virtual environment (VE) spatial memory task while they underwent fMRI. The task was programmed using UnrealEngine2 Runtime software (Epic, https://www.unrealengine.com) and participants were familiarized with all procedures before entering the MRI scanner, for details see Schuck et al. (2015). The VE displayed a grass plane surrounded by a circular, non-traversable stone wall with a diameter of 180 virtual meters (vm; 1 vm = 62.5 Unreal Units). Beyond the stone wall distal orientation cues, including multiple mountains, clouds, and the sun, were projected at infinite distance. Inside the arena a landmark was placed in the form of a traffic cone, see Figure 1. Participants were able to freely move around the arena. All movements were controlled using an MR-compatible joystick (NAtA Technology, Coquitlam, Canada) and exhibited constant speed. Right and left tilts of the joystick led to corresponding rotations of the player’s viewing direction. Forward and backward tilts controlled walking. A full crossing of the environment took approximately 15 seconds. Location and viewing direction of the player were recorded every 100ms.

**Figure 1:**
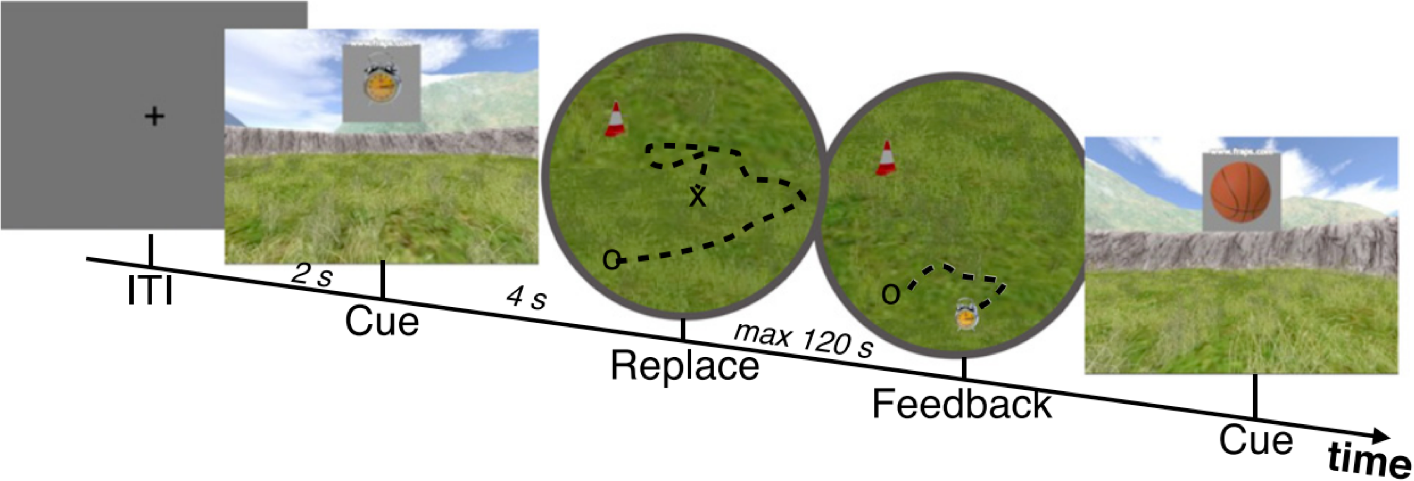
Trial structure of the VR task during feedback trials. After an object was cued it had to be placed at the remembered location. After replacing the object feedback was presented in the form of the true object location where it had to be picked up to start the next trial. Maximum time window for replacing and collecting was 120 s. Free movement during all parts of the feedback trials were used for further analysis. Figure adapted from Schuck et al. (2015).

Participants were first asked to encode the locations of objects that were shown within the arena. Afterwards, the participants’ main task was to navigate to the locations of these objects after a cue was displayed (see Figure 1; 5 objects, 6 trials per object, maximum time to relocate an object was 120s, for details see Schuck et al. (2015)). The analyses presented in this paper are solely concerned with directional signals independent of task condition. Thus, we considered all periods of fMRI recording that involved free navigation in a known environment. Encoding and transfer trials mentioned in the original publication were excluded since in encoding trials the environment was novel and movement was directed by cues and transfer trials involved changes to the environment that could potentially lead to direction remapping (e.g., Taube et al., 1990b).

### 2.3 Image acquisition

A 3 Tesla Siemens Magnetom Trio (Siemens, Erlangen, Germany) research-dedicated MRI scanner was used for MRI data acquisition. An MP-RAGE pulse sequence (1×1×1 *mm* voxels, TR = 2500 *ms*, TE = 4.77 *ms*, TI = 110 *ms*, acquisition matrix = 256 × 256 × 192, FOV = 256 *mm*, flip angle = 7°, 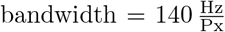) was used to collect T1-weighted structural images before and after the full task. Functional data was acquired using a T2*-weighted echo-planar imaging (EPI) pulse sequence (3 × 3 × 3 *mm* voxels, slice thickness = 2.5 *mm*, distance factor = 20%, TR = 2400 *ms*, TE = 30 *ms*, image matrix = 72 × 72, FOV = 216 *mm*, flip angle = 80°, 43 axial slices, GRAPPA parallel imaging, acceleration factor: 2, interleaved acquisition). Slices collected during the EPI sequence were rotated to approximately −30° relative to the anterior-posterior-commisure plane to reduce signal drop-out in areas of the MTL. The task was split into two functional runs, each taking between ten and 40 minutes depending on participant performance.

### 2.4 Image preprocessing

All imaging data were preprocessed and analyzed using SPM12. The pipeline for each subject consisted of spatial realignment, slice timing correction, coregistration to the anatomical scan and segmentation of the structural scan. Grey- and white-matter segmented anatomical images were used to create age-group specific MNI templates using SPM’s DARTEL (Ashburner & Friston, 2009) to avoid age effects resulting from normalization to a template based on younger adults. Anatomical ROIs were defined in MNI space using the Harvard-Oxford Cortical Atlas and the Talairach Atlas and afterwards transformed into the the subject’s individual functional space using the inverse of the participant-specific transformation matrix to the DARTEL template. All further analyses were conducted within-subject.

### 2.5 fMRI analyses

Participants could determine their orientation by tracking their own rotation and attending to the visually displayed distal orientation cues. The analysis was therefore focused on the following set of ROIs that had previously been related to (head) directional signals or visual processing: the retrosplenial complex (RSC), the subiculum, a joint hippocampus and entorhinal cortex ROI, the thalamus (Taube, 2007; Shine, Valdés-Herrera, Hegarty, & Wolbers, 2016), and primary visual cortex (V1). Although joystick movements resulted in relative direction changes which were independent of the travelled direction (a left tilt resulted in a left rotation relative to the direction before joystick movement), a ROI of the primary motor cortex (M1) was used to capture potentially spurious, motion-related effects on decoding and served as a baseline. Using a control ROI as our baseline also avoids issues inherent to performing population inference based on t-tests of decoding results against a numerical baseline (Allefeld, Görgen, & Haynes, 2016).

#### Univariate estimation of directional fMRI signals

Participants’ behavior was characterized by their walking direction. Walking direction could be derived from the angle of the vector connecting consecutively logged locations in the VE. Continuous navigation of each participant was segmented into separate periods (events) during which walking direction stayed within one of six, discrete, 60° bins for longer than 1 second. Stopping continuous movement or shifting walking direction beyond the border of a bin marked the end of an event. Viewing direction of the player was logged directly by the task program and matched the walking direction during forward walking. Backward walking periods were identified by marking periods during which viewing and walking direction differed by 180°(±20°). These events were excluded from the main analysis and considered separately (see below). The resulting direction events were then used to construct general linear models (GLMs) for univariate estimation of direction specific fMRI activation signals.

Since successive directions might be auto-correlated during free navigation (participants change more often from 30° to 60° than to 180°, etc.), performing a GLM on temporally auto-correlated fMRI signals can result in biased pattern similarities (Cai, Schuck, Pillow, & Niv, 2019). This effect can lead to spurious similarities between neural patterns of similar walking directions. We reduced this estimation bias by temporally and directionally separating adjacent events on the analysis level. Specifically, we separated odd and even numbered forward walking events and modelled them in two distinct GLMs. This separation of odd and even events ensured that events within the same GLM were separated by *at least* the minimal duration of another event (1 second) and resulted in an average of 5.1 TRs between two events, which corresponds to 12.25 s (SD=9.68). Such temporal separation exponentially reduced noise correlations between events, as can be illustrated by considering a 1-step autoregressive model of the form

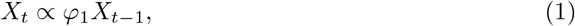

whereby *φ*_1_, known as the AR(1) coefficient, expresses the relation between the signal *X* at time *t* and the same signal during the previous measurement time-point *t* − 1 (constant and error terms are left out for simplicity). The relation between the signal *two* time steps apart can be found by substituting *X*_*t*−1_ in Eqn. (1) by its own auto-regression model, *X*_*t*−1_ *α φ*_1_*X*_*t*−2_, and is thus described by 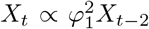. The now quadratic AR(1) term shows that the autocorrelation between the two measurements drops exponentially as a function of the number of ‘time steps’ between the measurements, i.e. the AR(1) coefficient of the signal recorded *p* time steps apart is an exponential function of the AR(1) coefficient *φ*_1_

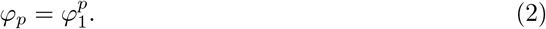

The value of *φ*_*p*_ comprises a signal component (similarity of directional representations) and a noise component (effects of previous noise components on following ones, e.g. caused by the slow nature of the hemodynamic response function). It therefore presents an *upper bound* of noise autocorrelation between consecutive events as some of the correlation might be due to similarities in directional representations. While the average AR(1) coefficient was .361 in RSC and .442 in V1, the correlations induced by temporal proximity between events in our GLMs were reduced to only .033 (SD=.046) and .063 (SD=.072) respectively. Note that these are average values over more detailed analyses which also revealed higher auto correlation in V1 for younger adults (for details see supplementary material section two). In addition to reducing temporal noise correlations, the separation of neighboring directions into more distant events ensured that temporally adjacent events mostly did not reflect neighboring directions, also reducing correlations among regressors (for details see supplementary material section three).

Directional GLM regressors were built to model data in each half run. Because the experiment contained two runs, events were split in four equal sets for each of the directions. This resulted in 24 direction regressors in total that were later used to perform cross-validated decoding. Direction regressors reflected onsets and duration of events as described above. The average event duration was 3.05 s (SD = 2.12 s). On average there were 114.98 events per subject (SD = 27.70). In addition, six run-specific motion and two run-wise intercept regressors were included, resulting in 38 regressors per GLM.

For an overview of the analysis pipeline see Figure 2.

**Figure 2:**
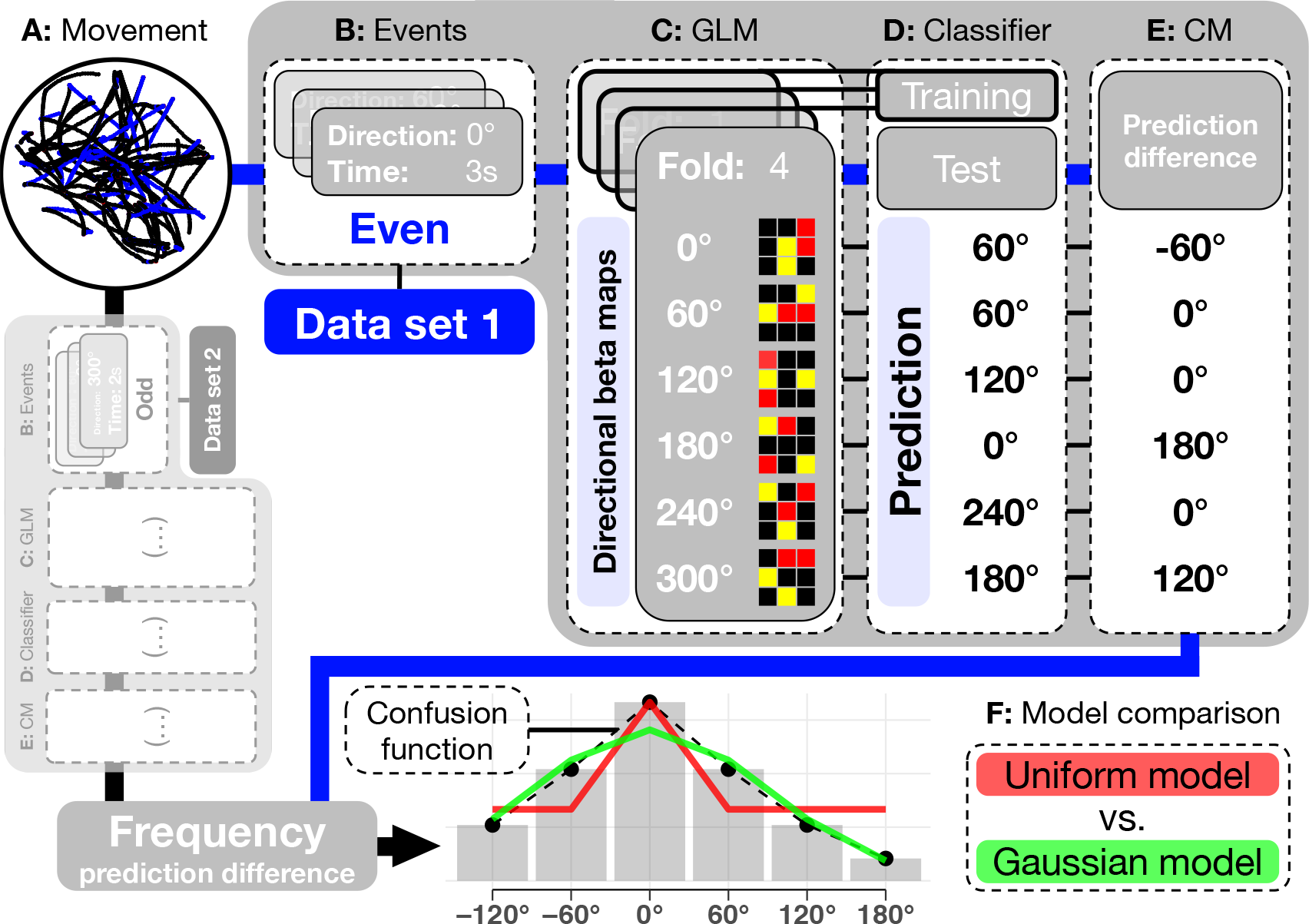
Schematic of analysis procedure. **A:** Individual navigation patterns during fMRI recording were separated into events corresponding to six possible angular walking directions. **B:** Odd (black) and even (blue) numbered events were analysed identically but as separate data sets to minimize confounds. **C:** Events entered a GLM yielding beta maps as representations of each walking direction in a four-fold structure. **D:** A classifier was trained on three of the four folds and predicted the walking direction for each beta map in the left out fold (exemplary numbers). **E:** Differences between predicted and true walking directions gave a direction invariant Confusion Matrix (CM). Confusion matrices were pooled over both data sets and normalized. **F:** Hypotheses concerning the predictive pattern of the classifier were tested by fitting two models: A uniform model assuming all false predictions are equally likely (*H*_0_; red) and a Gaussian model assuming errors to be less likely the more they diverge from the correct walking direction (*H*_1_; green). The Gaussian pattern should arise if the similarity of beta maps is a function of their angular difference, a prediction of a tuning function-like signal.

#### Classification of directional fMRI patterns

For all classification analyses, a multi-class linear support vector machine was trained on data from three folds and used to predict directions in the hold-out fold. Decoder training/testing was conducted using sciKit-learn (version 0.19.1, Pedregosa et al., 2011), nibabel (version 2.3.0, available at https://github.com/nipy/nibabel; Brett et al., 2018), and nilearn (version 0.4.1, available at https://github.com/nilearn; Abraham et al., 2014) packages in Python 3.6 (Python Software Foundation, version 3.6, available at http://www.python.org). Default settings (L2 penalty, penalty parameter *C* = 1, one-vs-rest multi-class strategy) and a maximum of 10^5^ iterations were used for classifier training.

Our main classification analyses were performed on direction-related beta maps and conducted separately for each GLM and ROI. Cross validated decoding results obtained from odd and even GLMs were averaged afterwards. In addition to testing the classifier on beta maps, we also applied it directly to data from single events. For each individual event, we calculated the precise average direction. This allowed us to relate classifier predictions to higher resolved direction labels of 10° per bin. Furthermore, the classifier was applied to backwards walking events. Because visual and walking direction diverged during backwards walking, this allowed us to quantify the influence of the visual scene on classification accuracy in different ROIs.

To test if classification accuracies exceeded chance level, accuracy levels in each ROI were compared to results from a permutation test (distribution of 1000 classification accuracies arising from training with randomly permuted labels) and to classification accuracy obtained in primary motor cortex (M1), using one-sided paired t-tests. P-values were Bonferroni-corrected for multiple comparisons across ROIs.

#### Influence of directional similarity on representational overlap of fMRI patterns

To test whether fMRI patterns that reflect similar walking directions are more overlapping than patterns associated with less similar walking directions, we analyzed the confusion matrix of the fMRI decoder. The confusion matrix reflects how often each category was decoded given a neural representation associated with each single category, e.g. how often did the classifier predict 120° although the walking direction was actually 60°, and etc. In a first step, we aligned the average proportions of classifier predictions around the true direction, and derived an average distribution of predictions around the true category, i.e. at − 120°, −60°, 0°, + 60°, + 120° and ±180°, relative to the target (averaged over folds and odd/even GLMs). This offered a *confusion function*, reflecting representation similarity/confusibility between two categories as a function of their angular difference (see Figure 2). We then quantified whether the confusion function reflected a tuning function by fitting a Gaussian bell curve to the data that peaks at the target direction, as done in electrophysiological animal research (Mazurek et al., 2014). The Gaussian curve is described by

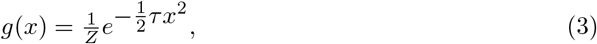

where *x* is a given direction relative to the target, *τ* reflects the precision of the Gaussian (1/*σ*^2^), and *Z* ensures normalization. This model had only one free parameter, precision *τ* that reflects the width of the tuning function. We compared this model to a null model that assumed evenly distributed off-target predictions independent of direction. According to this model, off-target predictions should be described by

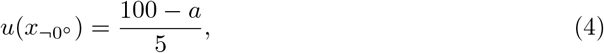

which uniformly distributes the percentage remaining after subtraction of the value at the target direction, *a*, a free parameter.

Both models thus had only one free parameter and were compared based on the sum of squared errors (SSE) between the model predictions and the confusion function. Decoding at the correct category (the center of the confusion function) was excluded from the curve fit analysis in order to make the tuning function analysis independent from overall decoding accuracy and avoid a bias towards the uniform model, where model prediction at the center is always matched to the data via the free parameter *a*. For each ROI, participants’ SSE differences between both models were entered into a one-sided t-test to test for a better fit of the Gaussian vs uniform model. We also derived a tuning function from the classifier predictions when applied to single forward or backward events by quantifying how often each of the 60 ° labels from the training set was predicted for each of the 10° bins in the test set.

We then compared the fitted precision parameters between age groups and ROIs, testing our hypothesis that directional information is encoded with higher precision in younger compared to older adults (one-sided t-test). Since the precision parameter was non-normally distributed for some cases, tests for group comparisons were chosen accordingly.

#### Effects of ROI and Age group on classification

To evaluate differences and interactions between ROIs and age groups, we used a model comparison between nested linear-mixed effects (LME) models. All models included a random intercept per participant. Fixed effects were entered in a stepwise inclusion approach: Model 1 included fixed effects of the intercept and the ROI, Model 2 included fixed effects of intercept, ROI and age group, and Model 3 included an additional interaction between ROI and age group. The three models were compared using a likelihood ratio test and followed up by post-hoc t-tests.

Using GLM derived beta maps for classification ensured fully balanced training sets. Yet, imbalances could still exist on the level of events from which regressors were constructed. To check for potential group differences in the number of direction events, a ‘class balance score’ was calculated that reflected the deviation of the event distribution from uniform (root mean squared error between the measured relative number of events belonging to each class and the corresponding uniform distribution). The number of events and balance score of each fold and subject entered a set of nested LME models similar to the ones described above. The models included intercept and age-group (Model 1). No differences between age groups in balance score were found (*χ*^2^(1) *≤* .745, *p* ≥ .388). Likewise, no difference between age groups in number of events were found (*χ*^2^(1) *≤* .150, *p* ≥ .698).

#### Differentiation of viewing and walking direction

The classifier was trained on forward walking events, during which viewing and walking direction were identical. During backwards walking, however, viewing and walking directions are opposed (180° shifted). Thus, the more a classifier depends on visual information, the more it will predict 180° shifted directions during backward walking. We therefore quantified the influence of visual information on decoding accuracy, as well as on the shape of the confusion function, by comparing classifier predictions for forward versus backwards walking events. Backwards walking events on average made up 26.8% (SD = 13.4%) of all events. Note that in both cases the classifier was trained on forward direction beta maps so the amount of backwards walking events did not influence the classifier’s predictions. Visual influence on direction signals was measured by calculating the relative differences in predictions at the target (0°) and opposed (180°) directions between the backward and the forward test set.

Additionally, we asked whether the influence of visual information was different in younger and older adults. This would hint towards a broader form of dedifferentiation compared to changes in the similarity structure of neural responses to a continuous stimulus. In each ROI, visual influence scores of both age groups were therefore compared using a Welch two sample t-test.

### 2.6 Behavioral analysis

A detailed analysis of the behavioral results can be found in Schuck et al. (2015). Briefly, location memory was quantified as the Euclidean distance between the remembered and true location during the feedback phase (distance score). Our analyses in the present paper focused on the relation between the Euclidean distance and measures for neural specificity. We therefore used Euclidean distance as the dependent variable in two linear models which contained the factors Age and one ROI specific measures of neural specificity (either decoding accuracy or Gaussian precision). All variables were z-scored before entering the linear model and analyses were conducted in R (version 3.6.1, R Development Core Team, 2011).

## 3 Results

### 3.1 Classification of walking direction

Classification accuracies for each ROI can be found in Figure 3. One-sided permutation tests (10^4^ iterations) indicated above-chance classification accuracy in the V1, RSC, and Subiculum (all *p*_*adj.*_ ≤ .006) but none of the other ROIs (*p*_*adj.*_ ≥ .054). Only decoding accuracy in the RSC- and V1 masks, however, exceeded classification level in M1 (both *t*(42) < 2.58, *p*_*adj*._ ≤ .033). While V1 classification can be expected to be based on visual signals, MRI sensitivity to directional signals in RSC is in line with other investigations (Shine et al., 2016). We therefore proceeded with only these two ROIs for which we had clear evidence we could measure directional signals in the present data set.

**Figure 3:**
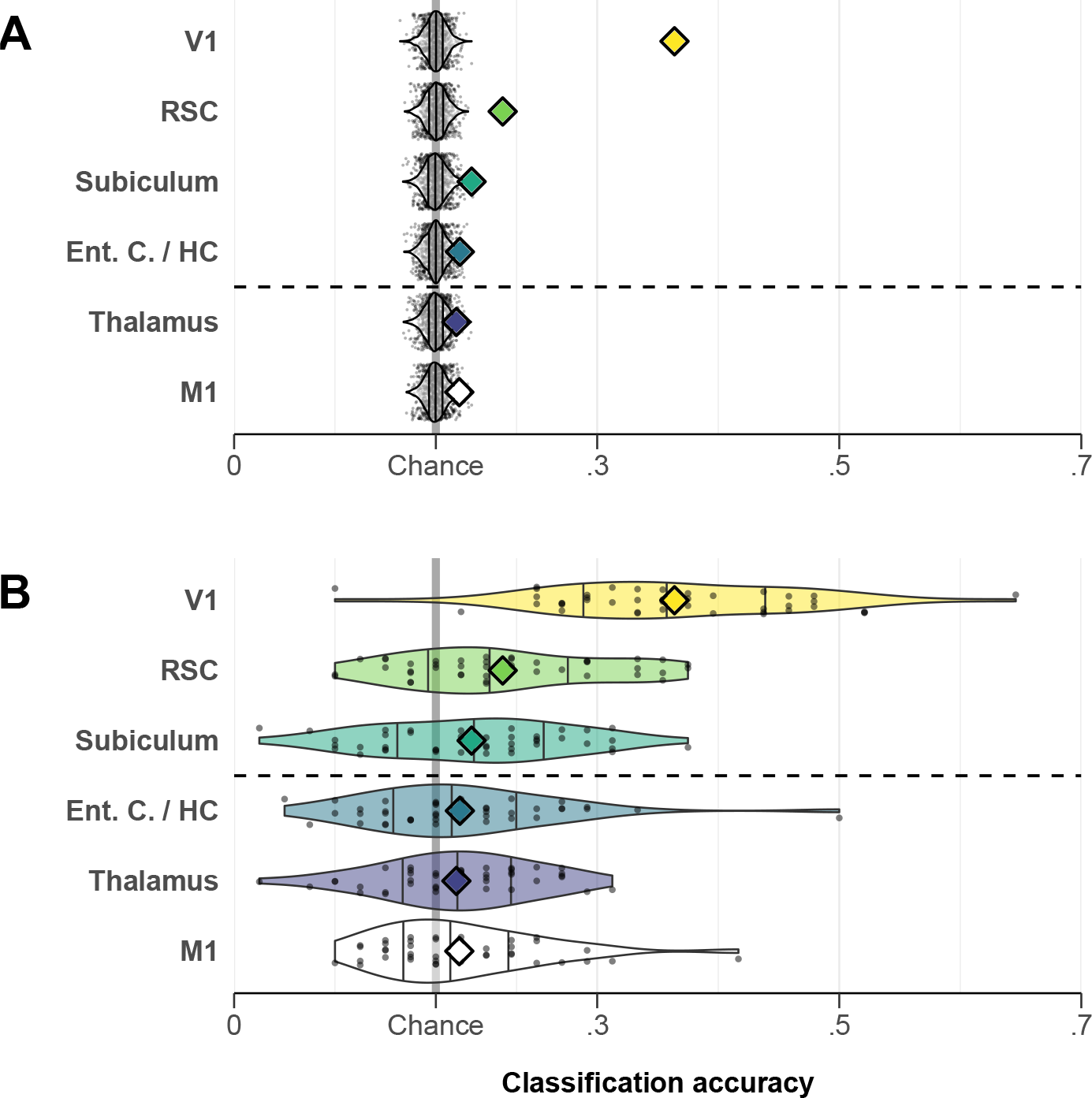
Decoder performance. **A:** Classification accuracy in each ROI (colored diamonds) compared to distribution arising from 10^4^ decoder runs with permuted labels (white violin plots, single values as black dots). Chance-level performance shown by grey line. ROIs above dashed line show significant above-chance accuracy measured by one-sided permutation tests and adjusted for multiple comparisons. **B:** Classification accuracies across investigated ROIs compared to M1. Single participant values shown as dots. Group means shown by color matching diamond. ROIs with significantly higher classification accuracies compared to M1 shown above dashed line.

Decoding accuracy tended to be higher in younger adults indicated by an increased model fit from including age group: *χ*^2^(1) = 10.90, *p* < .001) and was also higher in V1 than RSC (post-hoc t-test, *t*(41) = −8.72, *p*_*adj*._ < .001). The interaction between ROI and age did not further improve model fit (*χ*^2^(1) = 2.31, *p* = .072), indicating that age differences were not significantly different between ROIs. Post-hoc t-tests revealed a significant difference between age groups in V1 (*t*(75) = −3.66, *p*_*adj*._ < .001), but not the RSC (*t*(75) = −1.86, *p*_*adj*._ = .066).

### 3.2 Tuning function like representations of direction

To test differences in similarity structure of directional representations, we fitted a Gaussian and a uniform model to the classifier confusion patterns, as described above. A paired t-test of model SSEs across groups revealed that the Gaussian curve fitted the classifier confusions better than the opposing uniform model in both the RSC and V1 (all *t*(42) > 3.75, *p*_*adj*._ < .001). The Age × ROI interaction significantly improved model fit (*χ*^2^(1) = 4.305, *p* = .038). This reflects the fact that the Gaussian model fitted the data better in V1 compared to RSC in younger but not in older adults (post hoc tests: *t*(42) = −4.07, *p*_*adj*._ < .001 and *t*(42) = −.84, *p*_*adj*._ = .812, respectively). SSE comparisons can be found in Figure 4.

**Figure 4:**
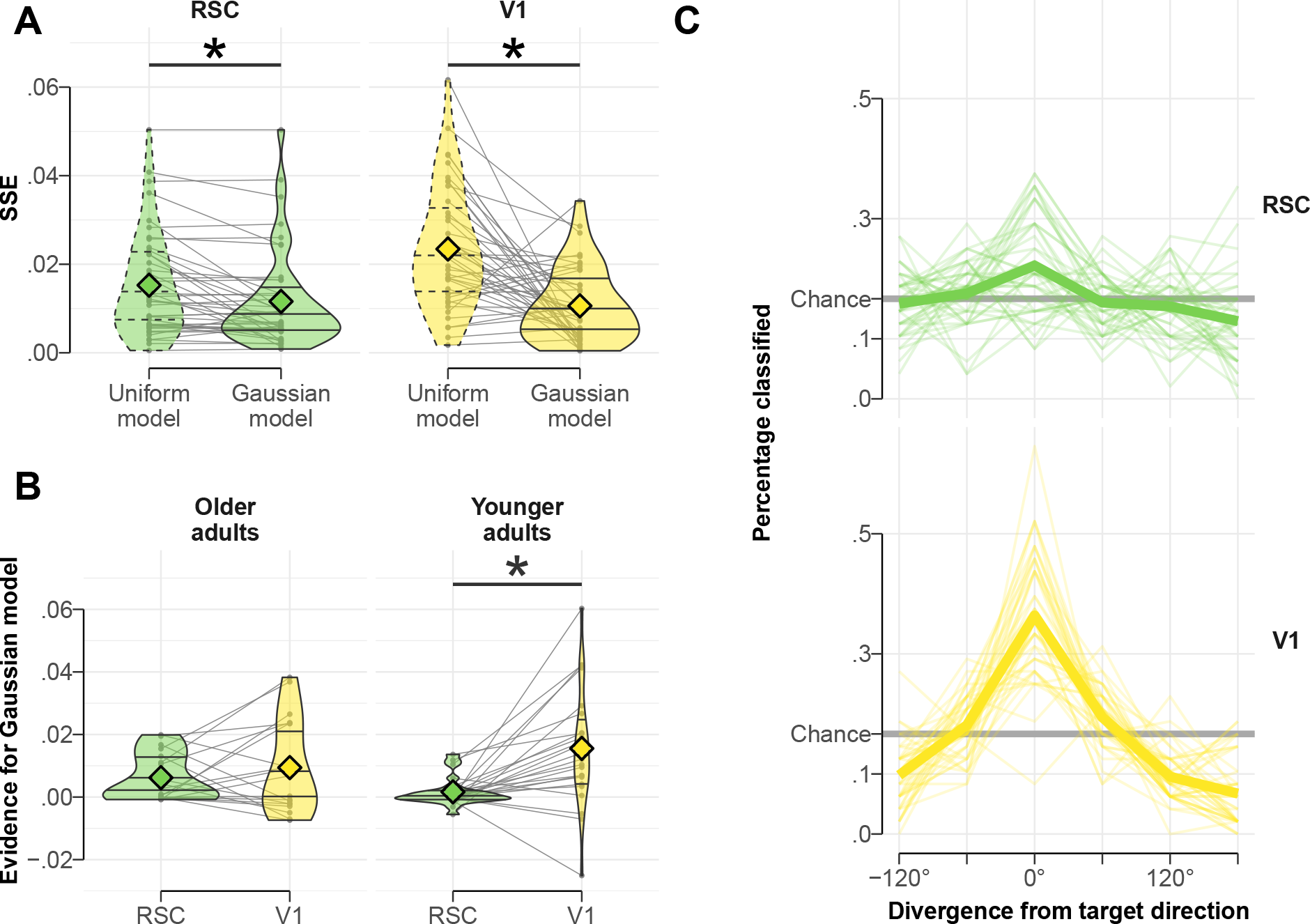
Quantification of tuning function-like signal **A:** Comparison between models fitted to confusion functions of RSC and V1 decoder. Depicted are within participant changes in SSE between models (violin plots as in Figure 3). Individual participant values for both models are connected by grey lines. Top dashed lines indicate significantly better fit of Gaussian model (within subject t-tests, one-sided, adjusted). **B:** Difference in model evidence when comparing RSC and V1 for both age groups. Evidence for Gaussian model is given by *SSE*_Uniform_ – SSE_Gaussian_, so values above 0 indicate a better fit of the Gaussian model. Dashed line indicates significant differences in a post-hoc t-test after correction for multiple comparisons. **C:** Depiction of confusion functions of RSC and V1 decoder. Participant specific confusion functions shown as thin lines. Thick line shows mean confusion function over all participants.

### 3.3 Differences in tuning width between age groups

Next, we investigated whether Gaussian precision differed between younger and older adults in either RSC or V1. Because normality was violated in at least one case (V1 precision in younger adults was non-normally distributed, Kolmogorov-Smirnov test, *D* = .515, *p* < .001), we used non-parametric Wilcoxon rank sum tests for these analyses. In V1, this test indicated significantly higher precision in younger compared to older adults (*W* = 138.5, *p*_*adj*._ = .029, one-sided). In the RSC, no such effect was found (*t*(40.98) =, *p*_*adj*._ = .917, Welch two sample t-test, one-sided). ROI-wise comparisons of precision and averaged confusion matrices in V1 for both age groups are shown in Figure 5.

**Figure 5:**
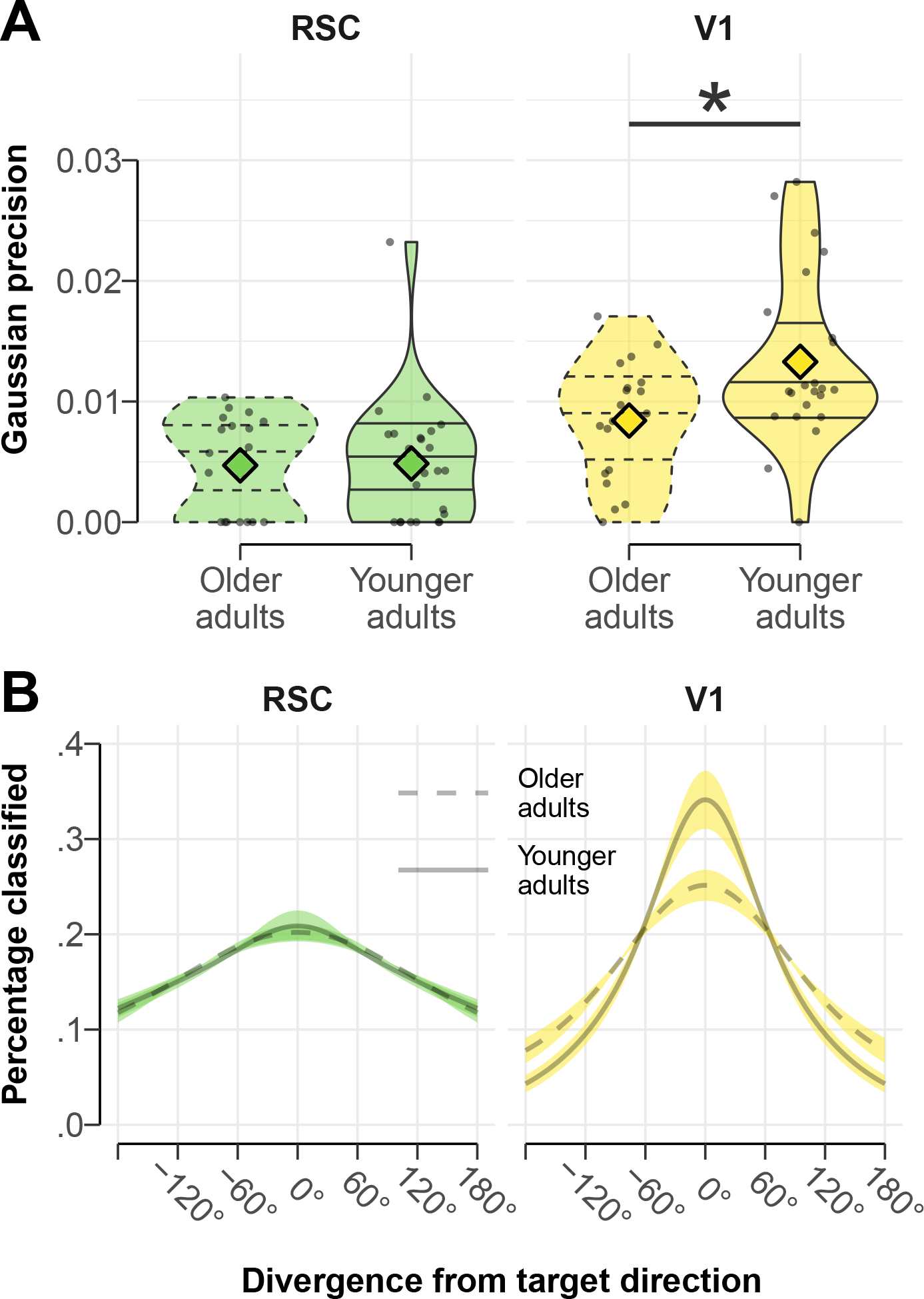
Tuning width of confusion function **A:** Group comparison of precision of Gaussian model fit to confusion function for RSC and V1, plots as in Figure. 3. Top dashed line indicates significantly higher precision in younger adults (one-sided t-test, adjusted). One high precision outlier (young participant) is not displayed in the V1 plot. **B:** Visualization of averaged best fitting Gaussian models of confusion functions for both age groups. Dotted lines and shaded area indicate standard error of the mean. Models were normalized to represent the percentage classified at the six measurement points of the confusion function.

To achieve higher resolution regarding the similarity of directional signals, and better support for our fitted models, we repeated the above analyses with classifier results when applied to the single event test set. Results of all analyses are shown in Figure 6. As expected, during forward walking events decoding accuracy was higher than in a permutation test in RSC as well as V1 (all *p*_*adj*._ < .001; see Figure 6A). Average high-resolution confusion functions can be found in Figure 6B. Applying the Gaussian and uniform models to the confusion functions indicated Gaussian like pattern similarites as expected in the RSC and V1 (*t*(43) *≤* −5.82, *p*_*adj*._ ≤ .001, paired t-tests of SSEs associated with each model; see Figure 6C). Similar to the classifier tested on beta maps, age-group differences in precision of the fitted Gaussians only showed a higher precision in younger adults compared to older adults in the V1 (*t*(29.95) = −3.47, *p*_*adj*._ = .001) but not in the RSC (*t*(35.82) = −.63, *p*_*adj*._ = .531, two sample t-tests, assumption of normality not violated; see Figure 6D).

**Figure 6:**
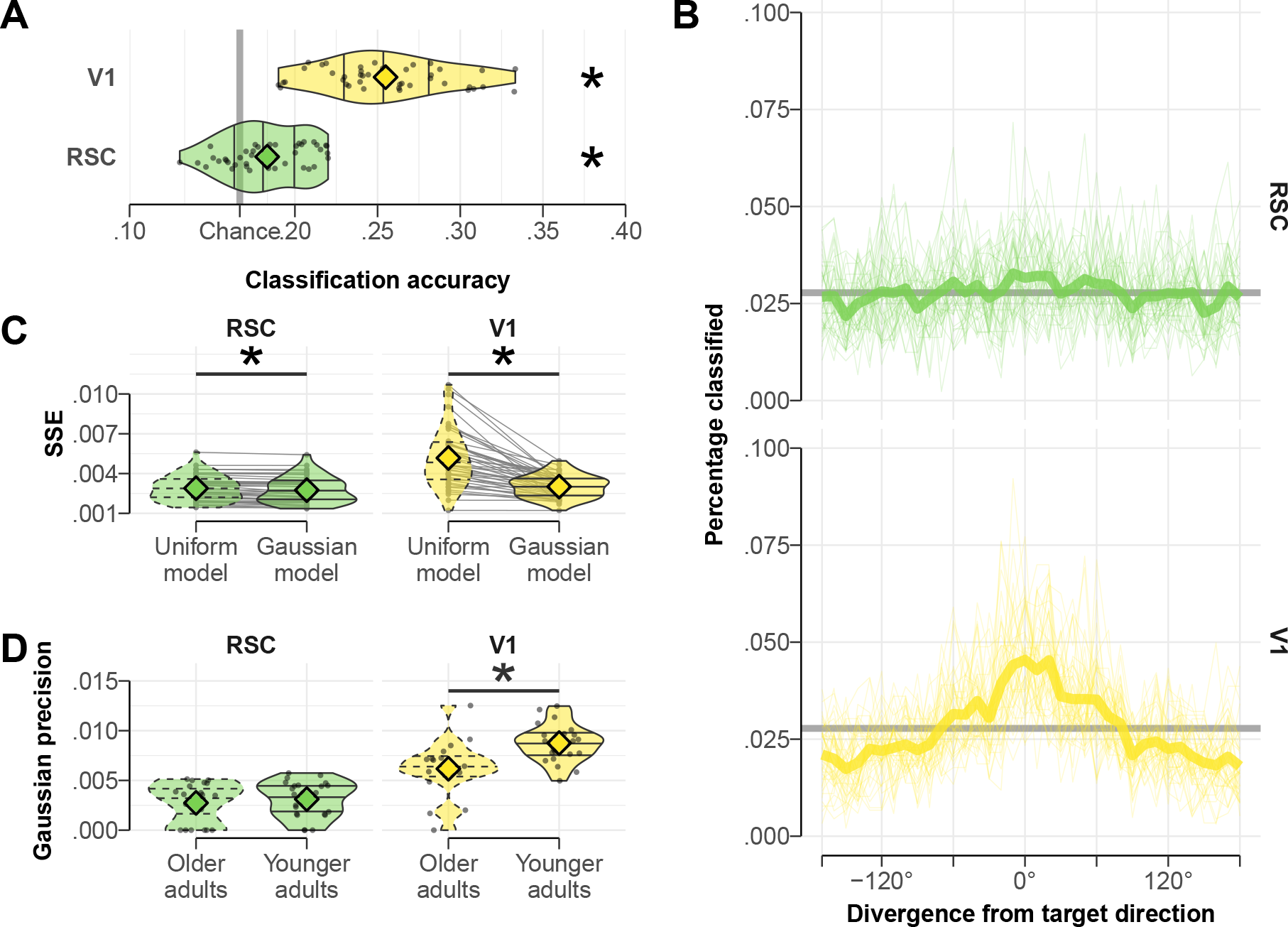
Analysis of decoders tested on single events. **A:** Classification accuracies of V1 and RSC decoders when tested on single events instead of beta maps. Depiction as in Figure 3B. Stars indicate significant above-chance classification accuracy given by a permutation test (10^4^ permutations, one-sided, adjusted). **B:** High resolution confusion functions of RSC and V1 decoder with a bin-width of 10°. Depicted as in Figure 4C. **C:** Comparison between models fitted to high resolution confusion functions of RSC and V1 decoder. Depicted as in Figure 4A. Top dashed lines indicate significantly better fit of Gaussian model (paired t-test, one-sided, adjusted). **D:** Group comparison of precision of Gaussian model fit to high resolution confusion function for RSC and V1. Plots displayed as in 5A. Top dashed line indicates significantly higher precision in younger adults (one-sided t-test, adjusted).

### 3.4 Influence of visual scene processing on decoding accuracy

Backwards walking events in the test set allowed us to investigate the influence of viewing direction on classification accuracy in each of the ROIs, since walking and viewing direction are opposite to each other. Visual influence on the directional signal was quantified as a decrease in (correct) predictions of walking direction combined with a simultaneous increase in 180° shifted predictions (in line with viewing direction) for backward relative to forward walking events. This measure of visual influence was then compared between ROIs.

A comparison of visual influence scores can be seen in Figure 7A. A paired t-test showed a significant difference of the visual influence between the V1 and the RSC ROI with lower visual influence in the RSC (*t*(42) = −7.15, *p* = .001), indicating qualitative differences in
the nature of the decoded representations in RSC versus V1.

**Figure 7:**
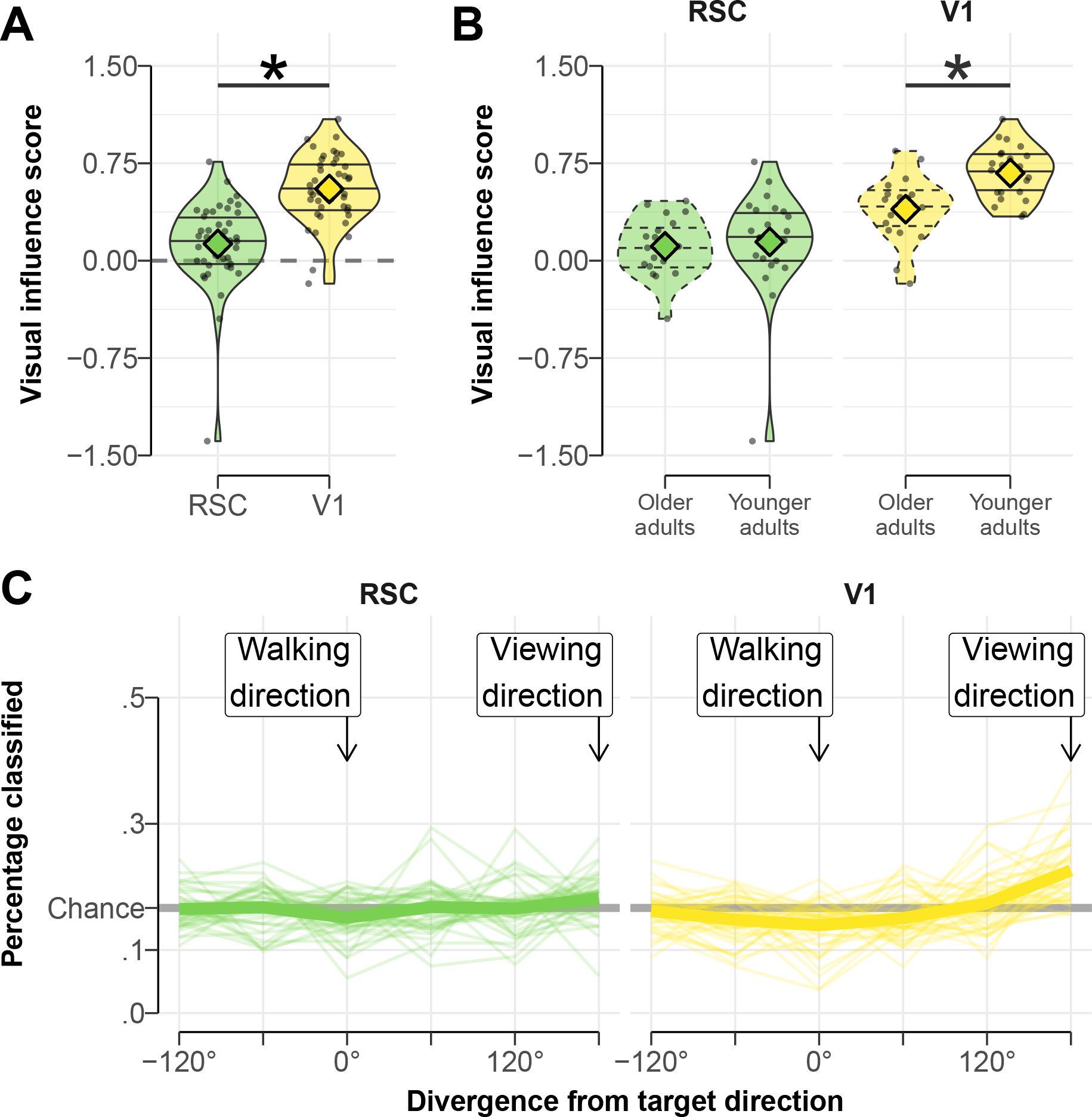
Influence of visual scene on direction prediction for RSC and V1. **A:** Comparison of visual influence score of RSC and V1 decoder. Plots as in Figure 3. Higher values above zero indicate a stronger tendency to predict the viewing direction while walking backwards. Dashed line indicates no visual influence. Top solid line and star show a significant difference in visual influence score between ROIs. **B:** Comparison of visual influence score between younger and older age group in each ROI. Solid line and star indicate significant comparison. **C:** Individual (thin lines) and average (heavy line) confusion function for RSC and V1 decoders tested on backwards walking events where walking and viewing direction are opposite to each other (180° divergence, indicated by labeled arrows). Functions peaking at 180° correspond to high visual influence.

We next asked whether the influence of visual information was different in younger and older adults, hinting at a broader form of dedifferentiation. Visual influence scores were lower in older adults compared to younger adults in V1 (*t*(33.28) = −3.95, *p*_*adj*._ < .001) but not RSC (*t*(37.51) = −.34, *p*_*adj*._ > .999). See Figure 7B for an age group comparison of visual influence scores. Confusion functions of the decoder trained on forward walking and tested on backward walking are shown in Figure 7C.

### 3.5 Behavioral results

We explored relations between task performance and measures for neural specificity in each ROI, predicting distance scores by age group and either decoding accuracy or Gaussian precision. Both measures of neural specificity stemming from a decoder trained and tested on directional beta maps. Predicting distance scores using the predictors age group and decoding accuracy in the RSC showed a negative relation with distance score independent of the age group, indicating worse task performance with less decoding accuracy (*r* = .172, *p* = .015, *r*_*younger*_ = −.542, *r*_*older*_ = −.279). Other linear models did not show any relation between distance score and measures of neural specificity that was independent of the age group. Relations between either decoding accuracy or Gaussian precision and distance score are shown in Figure 8A. and B., respectively.

**Figure 8:**
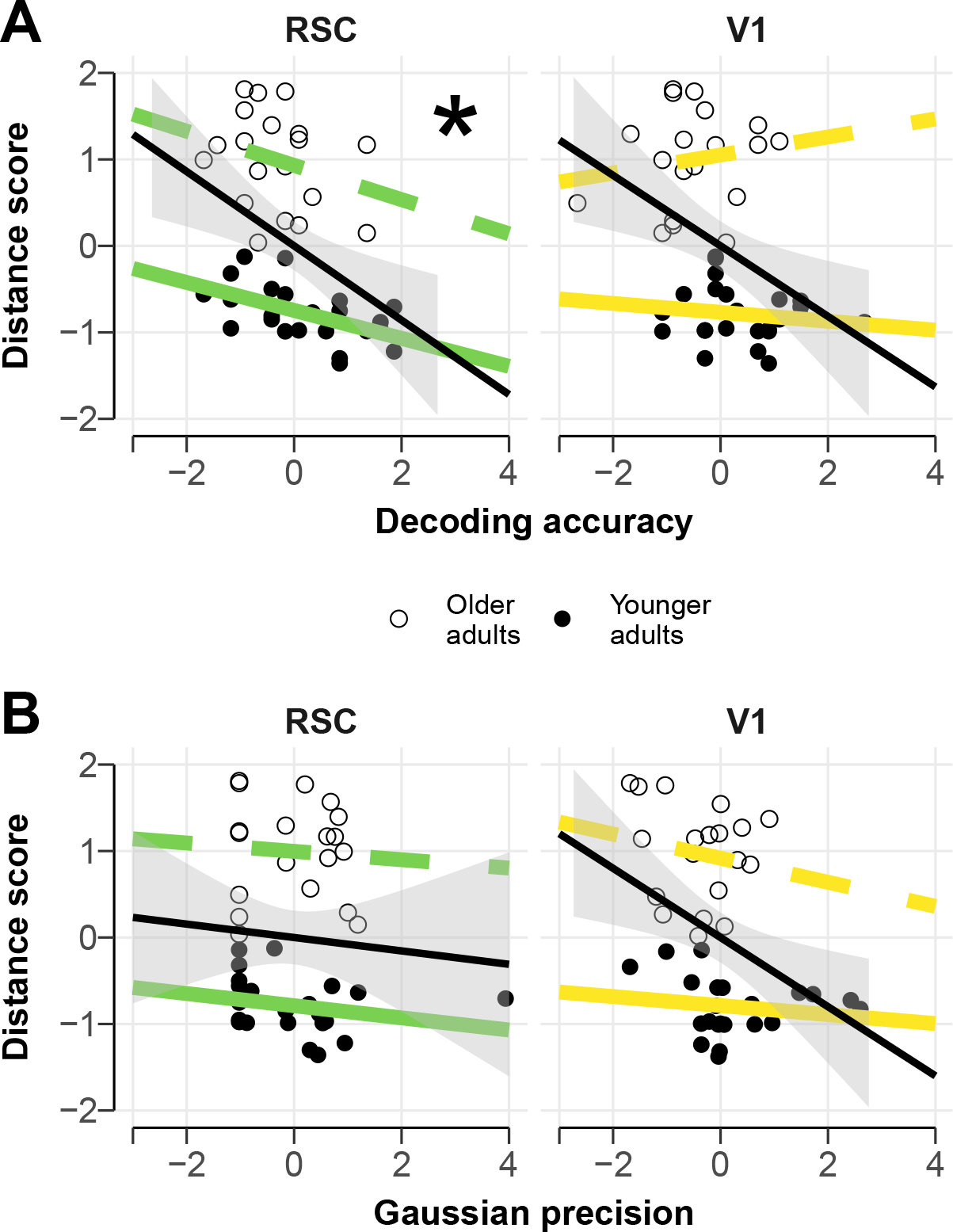
Relation between measures of neural specificity and task performance measured by distance score. Measures of neural specificity stem from a decoder trained and tested on directional beta maps. **A:** Relation between decoding accuracy and distance score. Younger adults shown by solid points and lines. Colored and black lines show correlation within and across age group, respectively. Star indicates a significant correlation independent of age group. **B:** Relation between Gaussian precision and distance score. Coloring identical to figure panel A.

## 4 Conclusions

In this study we used fMRI to investigate age-related changes in the specificity of direction-selective neural signals. More specifically, we asked a set of three hierarchically structured questions: whether it is possible to decode angular walking direction during free movement, if the similarity of neural patterns associated with these directions declines gradually with larger angular differences, as predicted by directional tuning functions, and whether older adults show broadened directional representations.

Our results revealed that directional information could be decoded from fMRI patterns in the RSC and V1, in line with previous investigations (Shine et al., 2016). Interestingly, age differences in decoding accuracy were found only in V1, but not RSC. Going beyond mere accuracy, we introduced a novel method that allowed us to characterize tuning function-like signals during decoder readout, while minimizing effects of autocorrelations. This analysis demonstrated that independent of overall classification accuracy decoder confusions in both ROIs were approximated best by a Gaussian tuning function – indicating a gradual decline of pattern similarity and following the predictions of a tuning-function like signal as found in animal research. Analyzing the width of the fMRI-level tuning function indicated broadened tuning of visual representations in V1 in older adults. In line with our predictions, this provides evidence for broader tuning functions in older adults as suggested by the neural broadening hypothesis (J. Park et al., 2012). Unexpectedly, no evidence for age differences in tuning width was found in RSC. Analyses for single trial events confirmed our results and showed that the Gaussian similarity structure persisted when directional signals were resolved at 10° instead of 60°. We also quantified the impact of visual information on direction decoding by analyzing backwards walking events and found that RSC signals were less contingent on the visual scene present than V1, as expected. Additionally, younger and older adults differed in the influence the visual scene had on the signal measured in V1, but not RSC.

To the best of our knowledge, this is the first study to investigate potential age-related changes in tuning functions defined over a continuous domain, rather than using discrete categories (J. Park et al., 2012; Koen et al., 2019). This is a notable distinction from previous research since mechanisms of age-related changes are likely different in these two cases: dedifferentiated responses within local circuits, which code the same continuous quantity, are related to changes in local inhibitory control, such as GABAergic interneurons (Leventhal et al., 2003); cross-areal dedifferentiation conceivably reflects a range of different mechanisms, including changes in long range connectivity or task strategies (Reuter-Lorenz & Lustig, 2005; Reuter-Lorenz & Cappell, 2008). Moreover, investigating continuous encoding of direction allowed us to test the claims made by the neural broadening hypothesis more directly: does a tuning function defined over continuous space change with age? Our findings in V1 converge with previous findings, but the apparent lack of evidence for age related dedifferentiation in RSC represents a notable deviation from previous findings and warrants further investigation.

While it is likely that the measured signal in the RSC or thalamus contains directional information influenced by head direction cells (Shine et al., 2016), effects in the V1 are most likely based on visual inputs drawn from a continuous visual scene. Our results suggesting neural broadening in the early visual system converge with findings in single cell recordings demonstrating wider tuning functions in senescent monkeys confronted with a visual stimulus of various orientations (Leventhal et al., 2003). While visual orientation signalling occurs earlier in the visual hierarchy than scene detection, it is possible that this process drives the present findings in V1 and suggests that the introduced method to investigate neural broadening might be sensitive to tuning curve related changes. Our results furthermore indicate that classifier confusions can pose as a tuning function proxy measure of a continuous variable, providing a novel measure for neural specificity beyond classification accuracy. To see if the findings are specific to the investigated domains, the method should also be applied to other continuous variables, e.g. the perception of motion and spatial frequency (Liang et al., 2010; Yang et al., 2008).

The observed relationship between a less specific directional signal in the RSC and larger errors in the placement of objects to memorized locations suggests that neural dedifferentiation might play a role in spatial memory performance. Since there was no group difference in classification accuracy in the RSC, it remains unclear if this process is connected specifically to the aging brain or rather describes a process which is happening throughout the adult lifespan (Rugg, 2016). This idea was also supported by a study by Koen et al. (2018) where the connection between neural dedifferentiation and memory performance was also age invariant.

The reason why no evidence for neural broadening and/or age-differences in directional signal specificity could be found in areas associated with a less visually dominated signal remains unclear. One possible explanation could be that during the VR task in the fMRI scanner no matching vestibular information is provided to the participant. The vestibular system has been identified as a possible source of internal noise during the process of path integration (Stangl, Kanitscheider, Riemer, Fiete, & Wolbers, 2018), a skill heavily relying on HD signal (McNaughton et al., 2006) and heavily influenced by older age (Adamo, Briceño, Sindone, Alexander, & Moffat, 2012). As this error source is eliminated by lying motionless during the task, age-differences might diminish. Furthermore, the resulting finding could have been limited by the resolution of directional categories. Smaller bins of directions when training the decoder would increase the resolution and accuracy of the investigated confusion functions. In order to exclude the possibility of neural broadening of directional signals in the human brain a similar approach with a more specialized paradigm should be conducted. It should however also be mentioned that, to the best of our knowledge, currently no evidence exists that directionally tuned cells are subject to neural broadening.

It is important to note that this paper presents a reanalysis of data collected during a task that was not specifically designed for the purpose of this study. It was therefore impossible to unambiguously disentangle visual from non-visual direction signals and travelled directions were not experimentally controlled. In consequence, visual input partially confounded directional signals, travelled directions were autocorrelated, some directional events occurred more frequently than others and some events that were unfit for directional analysis altogether (e.g. being idle and micro movements). We note that all these aspects are characteristics of navigation as it occurs in daily life and our analytical approach has shown how autocorrelations can be reduced and the amount of visual influence on neural representations can be characterized. Yet, the introduced limitations would be less severe in a tailored experiment which could increase analytic sensitivity. Another limitation regarding the generalization of these results includes the solely male participants. While this avoided effects based on participant’s sex the findings should not be generalized to female populations without further validation.

One important open question relates to the relation between our confusion function-based measure for neural specificity over continuous variables and changes in neurotransmitter systems. Contemporary models have linked neural dedifferentiation to less reliable or reduced DA-related signalling in the aging brain (Li & Rieckmann, 2014), a dominant aspect of the aging brain that is known to influence learning (e.g., Eppinger, Schuck, Nystrom, & Cohen, 2013) and memory (e.g., Schuck et al., 2013). The effect of changing DA levels in younger and older adults on neural specificity measured over a continuous variable could provide more detailed insights towards the mechanisms behind neural dedifferentiation and the role of DA in the aging brain. Moreover, understanding the role of GABA in this process is important given its known influence on neural broadening (Leventhal et al., 2003; Lalwani et al., 2019). Future studies employing neurotransmitter imaging, pharmacological interventions and genetic or pharmacogenetic approaches therefore promise to shed more light on age-related changes in ‘local’ tuning functions and cross domain dedifferentiation that characterize the human brain.

## Acknowledgement

This work was funded by a research group grant awarded to NWS by the Max Planck Society (M.TN.A.BILD0004).

We thank Douglas Garrett and Ulman Lindenberger for their helpful comments on the manuscript. Furthermore, we like to thank Lennart Wittkuhn, Nir Moneta, Samson Chien, Anika Löwe, and Ondřej Zíka for their remarks over the course of this project.

