## Supplementary Material for "Effects of Aging on the Encoding of Spatial Direction in the Human Brain"

### 1 Substituting missing events

In the case participants had no occurrences of certain directional events in one of the two data sets (even or oddly numbered events) the construction of a directional beta map for this direction would not have been possible. To obtain a beta map, the missing events in one GLM were substituted with events of the other data set. If an event was present in neither of the two data sets or this procedure had to be applied more than two times the participant was excluded. As a consequence, two older participants had to be excluded. This procedure had to be applied a total of 22 times across 15 participants (once in 8 participants, twice in 7 participants).

Another participant of the younger age group had to be excluded since not enough data was left after thresholding movement to at least 1 s.

### 2 Autocorrelative structure

AR(1) coefficients $\varphi_1$ were acquired for each participant and each voxel by correlating each time course with the same time course shifted backwards by one measurement point (TR). Voxelwise averaging yielded ROI-specific $\varphi_1$ values. $\varphi_1$ group averages for each ROI are shown in Table 1. Considering Eqn. (1) and (2) the remaining upper bound for noise auto correlation between two TRs can be estimated by $\varphi_1$ to the power of the elapsed number of TRs. By entering only even or odd numbered events into the same GLM the number of TRs elapsed between consecutive events inside the same GLM is increased. This effectively decreases the upper bound for noise auto correlation and changes patters of directional transitions.

Participant-specific average numbers of TRs elapsed between consecutive events were determined for each possible directional transition ($-120°$, $-60°$, $0°$, $60°$, $120°$, and $180°$) within the same GLM and averaged afterwards. Average values for each group can be found in Table 2. Utilizing participant specific values for $\varphi_1$ (different in each ROI), average elapsed TRs between consecutive events (different for each directional transition), and Eqn. (2) the remaining upper bound for noise auto correlation could be estimated for each ROI and directional transition and

| | ROI | | | | | |
|---|---|---|---|---|---|---|
| Age group | V1 | RSC | Subiculum | Ent. C. / HC | Thalamus | M1 |
| Older | .392 (.136) | .339 (.107) | .310 (.104) | .347 (.113) | .222 (.104) | .285 (.122) |
| Younger | .484 (.082) | .379 (.080) | .327 (.083) | .370 (.087) | .251 (.069) | .353 (.080) |

**Table 1:** Average $\varphi_1$ values and standard deviation (in brackets) for age groups and each investigated ROI.

are shown in Figure 1.

| | Directional transition | | | | | |
|---|---|---|---|---|---|---|
| Age group | $-120°$ | $-60°$ | $0°$ | $60°$ | $120°$ | $180°$ |
| Older | 5.07 (4.39) | 5.98 (5.42) | 6.12 (5.72) | 6.19 (4.76) | 6.27 (4.96) | 7.41 (4.63) |
| Younger | 4.69 (2.86) | 4.68 (3.04) | 3.62 (2.76) | 3.99 (2.79) | 4.32 (2.82) | 5.10 (2.70) |

**Table 2:** Mean and standard deviation (in brackets) of average TRs between consecutive events. Resolved by directional transition and age group.

### 3 Disrupting structure of directional shifts

In addition to a reduction in auto correlation, separating events in two data sets (odd and even events) changes the distribution of directional transitions between consecutive events. Effects of separating events on directional transitions are displayed in Figure 2.

Since all participants freely navigated the environment, directional transitions happen gradually (e.g. slightly turning). Consequently, the most probable directional shift was between neighbouring directions ($\pm 60°$) if consecutive events were not separated. When separating consecutive events into two data sets (odd and even events) this transition structure was disrupted.

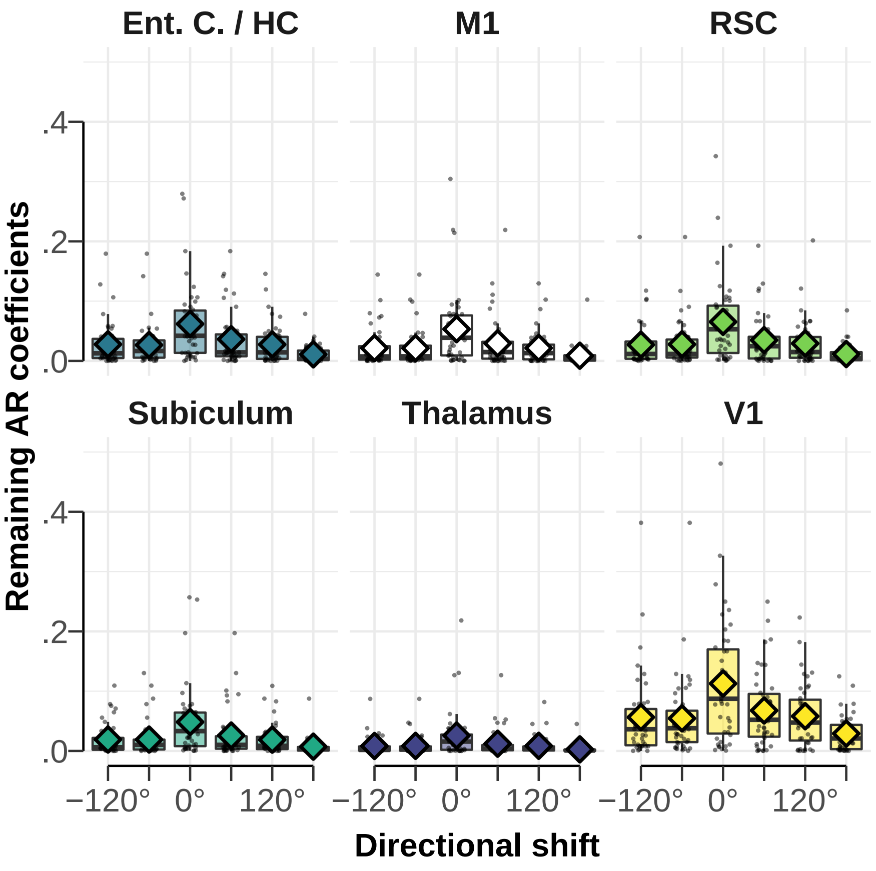


**Figure 1:** Estimation of remaining upper bound for noise auto correlation after considering elapsed TRs based on Eqn. (2). Individual remaining auto correlation after separating odd and even events are shown by black dots. Mean shown by diamonds. Because the number of elapsed TRs is different for each possible directional transition and $\varphi_1$ values are different for each ROI the estimates are resolved by ROI and directional transitions.

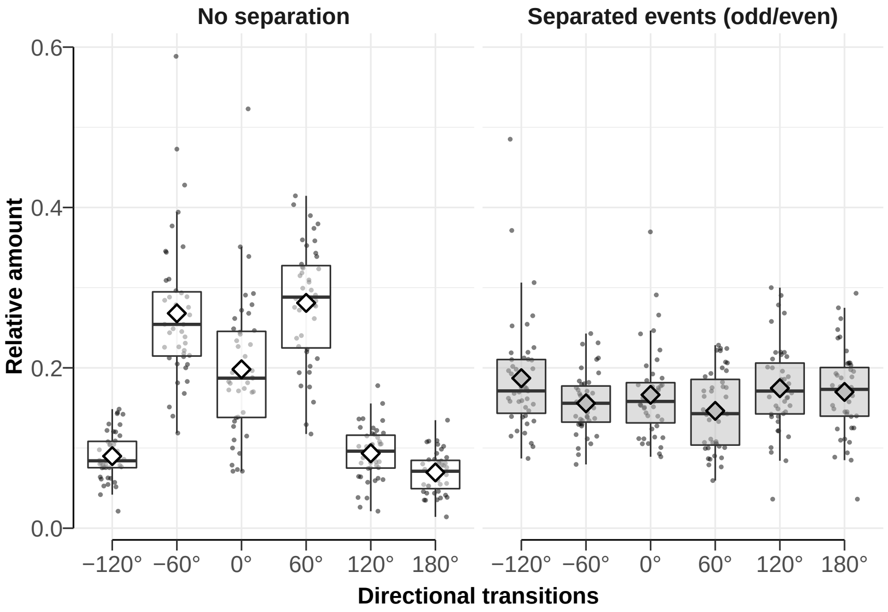


**Figure 2:** Influence of separating odd and even events into two data sets on the relative amount of directional transitions between consecutive events. Individual relative amount of directional transitions for each participant shown by black dots. Mean indicated by diamonds. When directional events are not separated the most prominent directional transitions are neighboured directions ($\pm 60°$). Applying said separation disrupts this transition structure.
